# Candidate gene mining of a fragmented *Tipularia discolor* Pursh (Nutt.) genome assembly reveals genes putatively associated with specialized metabolite production

**DOI:** 10.64898/2026.11.16.767051

**Authors:** Jonathan I. Watkinson

## Abstract

Orchids are a large and diverse group of plants with unique adaptations that have enabled the utilization of many different environmental habitats across almost all continents. These adaptations are mediated by an as-yet-underexplored genetic resource. Nearly all sequenced orchid genomes are of Asian origin and represent less than 0.1% of all orchid species, limiting the ability to explore and mine the gene space of this complex plant family. Greater representation of orchids in genome databases could provide important insights into the genes responsible for adaptive phenotypes, which could be useful for targeted crop improvement and in orchid conservation efforts. To begin to expand the palette of orchid genomes, an effort was undertaken to sequence the genome of a North American terrestrial orchid, *Tipularia discolor*. K-mer analysis estimated a haploid genome size of approximately 1.7 Gb, with a high proportion of repetitive sequences. Although a complete genome was not obtained, the implementation of a conservative gene mining approach and the generation of a candidate gene mining pipeline led to the identification of 68 consensus contig objects containing whole or partial genes related to sucrose, trehalose, terpenoid, and phenylpropanoid metabolism. Of note, two bibenzyl synthase homologs were identified with high confidence and clustered with characterized orchid bibenzyl synthases in a CHS/BBS phylogeny, indicating that *T. discolor*, like its Asian and South American relatives, has the capacity to produce this distinct class of phenylpropanoids. Thus, despite substantial fragmentation, the *T. discolor* assembly contains sufficient gene space representation for targeted candidate gene discovery. This works lays the foundation for functional characterization of selected candidates and development of a more complete genome assembly for comparative genomic studies.

## Introduction

Orchids are considered to be the largest and most diverse plant family and are known to occupy numerous environmental niches on almost every continent. This global success is due to various adaptation mechanisms, including a unique interaction with mycorrhizal fungi (Rasmussen and Rasmussen, 2008; Rasmussen et al., 2015), utilization of the epiphytic realm (Collobert et al., 2023), dynamic pollination syndromes (Ackerman et al., 2023; Kuriakose, 2026), and multiple examples of CAM photosynthetic pathways (Nivedha et al., 2026; Zeng et al., 2026). These adaptations comprise an underexplored genetic resource with the potential to provide genes relevant to directed plant improvement programs. To date, the genomes of approximately 30 species of orchid have been sequenced (Chen et al., 2022), most of which are of Asian origin with about half in the genus *Dendrobium* (Chen et al., 2025). These efforts reveal substantial differences in genome size and provide insights into genes governing floral morphology, another distinctive attribute of orchids (Chung et al., 2022). These sequencing efforts, however, encompass less than 0.1% of all orchids, suggesting that much still needs to be uncovered to answer questions related to orchid evolution and the genetic diversity underlying adaptation (Bailashaki et al., 2020). For example, comparative genomics between epiphytic and terrestrial species may yield insights into genes that enable different habitat choices (Collobert et al., 2023). Expanding the repertoire of orchid genome sequences will also provide an opportunity to mine the gene space for candidates that can be used in crop improvement (Tiwari et al., 2024) and generate insights into orchid biology that can be used to enhance conservation programs for this highly-threatened group of plants (e.g., Hartvig et al., 2026).

*Tipularia discolor* Pursh (Nutt.) is a terrestrial, North American orchid species that is widely distributed along the Eastern half of the United States. It is a member of the *Epidendroideae* subfamily, a clade largely dominated by epiphytic plants. Its habitat includes low, damp areas as well as dry slopes and it can be found in most woodlands in the mountains of southwest Virginia. Thus, insights from genome analysis may provide context for how orchids evolved to utilize the terrestrial habitat. In fall, a single leaf arises from the current season’s tuber, which is attached to the previous season’s tubers by a short rhizome. Roots emerge from the most recent tubers. After the leaf senesces in late spring, a single inflorescence arises from the previous season’s tuber with flowering occurring in late July into August. This unusual life cycle, with photosynthetic ability confined to the cold, low-light winter months, raises the possibility of a substantial contribution from mycorrhizal or other endophytic fungi. Intriguingly, the related genus *Corallorhiza* is achlorophyllous and strongly dependent on fungal partners.

Furthermore, a recent study has shown that habitat disturbance influences the community structure of *T. discolor* fungal endophytes, strengthening the argument that the mechanisms that underlie these interactions should be explored (Watkinson, 2025).

One of the unique aspects of orchid growth and development is a reliance on mycorrhizal fungi for germination and sustained growth (Alghamdi, 2019; Li et al., 2021; Rasmussen et al., 2015). Recent research highlights a broader role for fungi beyond mycorrhizal symbiosis, with numerous reports of communities of endophytic fungi also associated with orchid roots (Hu et al., 2024; Jeldes-Cajas et al., 2026; Ma et al., 2015; Novotná et al., 2018; Watkinson, 2025; Yu et al., 2015). Speculation about the role of these endophytic fungi ranges from contributing to increased growth through fungal-mediated nutrient uptake to enhancing survivability by contributing antimicrobial activity (Bhatti et al., 2022; Moreno-Peimbert et al., 2026), however the genetic mechanisms underlying orchid-fungal relations remain poorly understood. The exchange and regulation of specialized metabolites may represent one mechanism by which these plant–fungal interactions are mediated.

Orchids are also known for interactions with pollinators (Ackerman et al., 2023). This interaction can be highly specific and rely on both visual and olfactory cues. *T. discolor* was documented to be visited by noctuid moths in the late 1970’s, suggesting that olfactory cues may contribute to pollinator attraction (Natale, 2025; Whigham and McWethy, 1980). This research needs to be updated, particularly in light of climate shifts and the increased disturbance of habitats. However, the pollination strategy does indicate that *T. discolor* has the potential to produce an array of volatile compounds that contribute to reproductive success.

Many volatile compounds associated with pollinator attraction are derived from specialized metabolic pathways that also contribute to plant interactions with other organisms. In addition, evidence that endophytes can influence reproductive success suggests potential biochemical connections between above- and below-ground interactions (Schatz et al., 2010; Wolfe et al., 2005). Many questions could thus be addressed by examining the *T. discolor* gene space for candidate genes that mediate mycorrhizal formation, tolerance of endophytes, mechanisms that distinguish between beneficial and pathogenic fungi, and how fungi regulate gene expression in the host plant, which would require greater knowledge of the genes and genomic structure of the host plant.

To begin to address some of these questions, an effort was undertaken to generate a draft genome of *T. discolor* using Illumina paired-end short-read sequencing. While the overall objective of establishing a more complete genome assembly remains ongoing, the fragmented assembly contained sufficient representation of gene space to support targeted candidate-gene discovery. Using a conservative, protein-guided mining strategy, candidates associated with selected carbon metabolism and specialized metabolic pathways were identified. This approach provides an initial glimpse into the biochemical potential of *T. discolor* and establishes candidates for directed functional studies of its interactions with fungi and other organisms.

These genomic resources also provide a foundation for future sequencing and assembly efforts and comparative studies of orchid growth, development, physiology, and evolution.

## Methods

### Sequencing assembly

Leaves of *T. discolor* were collected in March 2026, and genomic DNA was extracted using the QIAGEN DNeasy Plant Mini Kit. Approximately 2 µg of DNA (260/280 = 1.83, with limited fragmentation based on gel analysis) was submitted to Novagene (Novagene USA, Sacramento, CA) for paired-end sequencing on the Illumina NovaSeq X Plus platform. Novagene quality-control metrics indicated approximately 333 million paired-end reads, an estimated sequencing error rate of 0.01%, and a GC content of approximately 38% (Table 1).

**Table 1.** Summary of Novogene Illumina Sequencing Data.

| Metric | Value |
|---|---|
| Platform | NovaSeq X Plus |
| Filtered Reads | 333 million paired-end |
| Read length | 150 bp |
| Total bases | ~100Gb |
| Estimated depth | ~60x |
| Estimated error (Novogene QC) | 0.01% |
| %GC | 38% |

Illumina reads were analyzed by 31-mer counting with Jellyfish (version 2.2.10; Marçais and Kingsford, 2011). The resulting k-mer frequency histogram was analyzed using GenomeScope 2.0 to estimate genome size, heterozygosity, repetitive content, and read error.

Genome assembly was performed using Assembly By Short Sequences (ABySS; (version 2.3.1;Jackman et al., 2017) and then using the Maryland Super-Read Celera Assembler (MaSuRCA version 4.1.4; Zimin et al., 2017). Because the MaSuRCA assembly did not yield populated conventional scaffold or contig FASTA files, consensus contig objects (CCOs) recovered from the completed Celera Assembler terminator stage were used for downstream gene-content analysis following an initial screen using BUSCO (Benchmarking Universal Single-Copy Orthologs) against the Embryophyta dataset (Simão et al., 2015).

### Candidate gene analysis

Reference protein sets representing selected genes and gene families involved in carbon metabolism, signaling, and specialized metabolism were assembled from UniProt (July 2026), with preference given to experimentally-characterized proteins where available. Reference sets, which varied in size and taxonomic breadth, were designed to provide protein-guided searches. The protein sequences used in each search category are provided in Supplementary Table S1.

Protein references were aligned to the MaSuRCA CCO collection using miniprot (version 0.13; Li, 2023), which performs protein-to-genome alignment while accounting for introns and frameshifts. Candidate CCOs were initially prioritized based on convergence of multiple reference proteins on the same CCO, alignment score, amino-acid identity, and coverage of the reference protein. Where multiple references converged on the same candidate region, the highest-quality reference to model alignment was retained for subsequent assessment. For selected genes in which fragmentation was suspected to preclude identification of candidates using full-length references, targeted searches were performed using selected portions of the reference proteins. The derived query sequences are provided in Supplementary Data File S2. Candidate CCOs identified through these focused searches were evaluated using the same validation criteria as candidates recovered in the initial screens.

Predicted protein models corresponding to identified candidate CCOs were extracted from miniprot-derived models or the corresponding focused searches (Supplementary Data File S3). Protein models were searched against the Pfam-A domain database (Payson-Lafosse, 2025) using HMMER (version 3.4; Eddy, 2011). Proteins were considered validated when they contained the expected domains with high significance scores and had the correct domain architecture. Significant matches were retained (Table 5). To provide additional support, conserved sequence motifs were also examined in some subsets of candidate proteins. Candidate annotations were therefore based on combined protein alignment, domain, and motif evidence rather than sequence similarity alone. Confidence in candidate identity was assigned using the combined evidence from reference coverage, miniprot alignment score, amino-acid identity, Pfam domain support, and where applicable, conserved motifs. Candidates were classified as very high confidence when protein alignment, expected domain architecture, and sequence completeness were strongly supported; high confidence when alignment and domain evidence were strong but the model was less complete; moderate confidence when homology and/or domain evidence supported the assignment but substantial sequence was missing or disrupted; and low confidence when only partial or comparatively weak alignment and domain evidence were available.

### Phylogenetic analysis

To further evaluate the identity of the putative *T. discolor* bibenzyl synthases, ClustalW was used to align protein sequences deduced from the three predicted type III polyketide synthase genes with reference CHS and BBS sequences from diverse monocot and dicots, including characterized BBS proteins from several orchid species. Stilbene synthases were added from grape and pine (see Supplementary Table S5). A phylogenetic tree was inferred using FastTree. The resulting Newick tree was rooted using a *Streptomyces coelicolor* type III polyketide synthase (WP_011027653) as an outgroup and plotted as a cladogram using the ape package in R (Paradis and Schleip, 2019).

### Computing

All computing was performed on the Virginia Tech Advanced Research Computing (ARC) cluster. Scripts used to perform analysis were written with the aid of ChatGPT.

### Accession numbers

This Genome has been deposited at DDBJ/ENA/GenBank under the accession JCDMZT000000000. The version described in this paper is version JCDMZT010000000. The BioProject number is PRJNA1521864.

### Supplementary material

Supplementary Table S1 contains the protein reference sequences used for candidate-gene mining. Supplementary Data File S2 contains derived protein query sequences used for targeted searches of selected fragmented candidates. Supplementary Data File S3 contains predicted protein models corresponding to the 68 candidate genes reported in Table 5 and used for downstream domain, motif, and functional assessment. Supplementary Data File S4 contains the nucleotide sequences of the 68 candidate CCOs reported in Table 5 (note that one CCO is not present in the sequence data submitted to NCBI as it was below the 1kb cutoff. Its retention in the supplemental data reflects the confidence in the putative annotation).

SupplementaryTable S5 lists the accession numbers for the proteins used in the phylogenetic tree analysis.

## Results

### Genome Assembly and Characterization

Orchids have unique biological adaptations that have the potential to provide useful traits to targeted crop improvement programs. In an effort to identify and characterize genes associated with these adaptive mechanisms, sequencing of the *T. discolor* genome was undertaken. In order to estimate the size of the genome, Novogene generated paired-end reads were subjected to Jellyfish K-mer analysis. GenomeScope analysis of the Jellyfish k-mer output estimated the *T. discolor* haploid genome at approximately 1.67–1.70 Gb (Table 2), comparable in size to other published orchid genomes (e.g., *Phalaenopsis*, 1.0 Gb; *Dendrobium*, 1.2 Gb; Cai et al., 2012; Zhang et al., 2016). The genome was estimated to contain a high proportion of repetitive sequence (55–56%) similar to that reported for Phalaenopsis but less than the 78% reported for Dendrobium (Cai et al., 2012; Zhang et al., 2016). Estimated heterozygosity ranged from 0.75% to 13.2%, depending on model parameters (Table 2), and the estimated read error rate was 0.85%.

**Table 2.** GenomeScope estimates of genomic parameters based Kmer analysis of 333 million Illumina PE reads.

| Metric | Value (Min) | Value (Max) |
|---|---|---|
| Homozygous | 86.73%s | 99.25% |
| Heterozygous | 0.745% | 13.27% |
| Genome Haploid length | 1,665,392,427 bp | 1,702,554,420 bp |
| Genome Repeat length | 934,809,663 bp | 955,669,245 bp |
| Genome Unique length | 760,582,764 bp | 746,885,175 bp |
| Model Fit | 45.49% | 85.52% |
| Read error rate | 0.855% | 0.855% |

Initial assembly with ABySS produced 9.61 Gb distributed among approximately 74.5 million sequences, with an N50 of only 129 bp, indicating extreme fragmentation (Table 3). Assembly was therefore repeated using MaSuRCA. Although the MaSuRCA pipeline did not produce populated conventional scaffold or contig FASTA files, approximately 5.1 Gb of consensus contig-object (CCO) sequence was recovered and used for downstream gene-content analysis (Table 3). The CCO collection remained highly fragmented; however, approximately 3.50 Gb (68.5%) of assembled sequence was contained in CCOs ≥1 kb, with a substantial tail of longer CCOs (Table 4).

**Table 3.** Comparison of assemblies from ABySS and MaSuRCA.

| Metric | ABySS | MaSuRCA |
|---|---|---|
| Assembly size | 9.61 Gb | 5.11 Gb |
| Sequences (contig objects) | 74,545,667 | 5,569,448 |
| Largest Contig | 15,678 bp | 76,266 bp |
| N50 | 129bp | 2333 bp |

**Table 4.** Consensus Contig Object size distribution.

| Size | Number of sequences | Bases represented |
|---|---|---|
| <200bp* | 1,024,928 | 0.17 Gb |
| 200-999bp* | 3,451,017 | 1.44 Gb |
| 1-1.99kb | 539,037 | 0.76 Gb |
| 2-4.99kb | 378,634 | 1.16 Gb |
| 5-9.99kb | 127,767 | 0.87 Gb |
| 10-20kb | 42,736 | 0.57 Gb |
| >20kb | 5329 | 0.13 Gb |

\*CCOs of less than 1kb were removed prior to submission to NCBI.  Sequences greater than the <1kb cutoff were deposited at DDBJ/ENA/GenBank under the accession JCDMZT000000000.

In order to validate the CCO collection and determine whether it captured some of the coding gene space, BUSCO analysis was performed. This analysis showed that 62.1% (1001 of 1614) of conserved Embryophyta orthologs were represented in the MaSuRCA CCO collection (Fig. 1), with 26% represented as complete genes and 36.1% as partial genes. Of the complete genes identified by BUSCO, 29 (6.9%) were classified as duplicated. Examination of duplicated sequences showed that most shared less than 90% identity, suggesting that these were more likely to represent related gene copies than near-identical assembly duplicates. Of the two pairs exceeding 90% identity, only one appeared to be consistent with assembly redundancy, with nearly the entire 1,488 bp CCO contained within a 6,424 bp CCO. The other involved much larger CCOs sharing only a limited region of approximately 91% identity. These results indicate that, despite substantial fragmentation, the assembly contained sufficient representation of conserved gene space to support targeted candidate-gene discovery.

**Figure 1.**
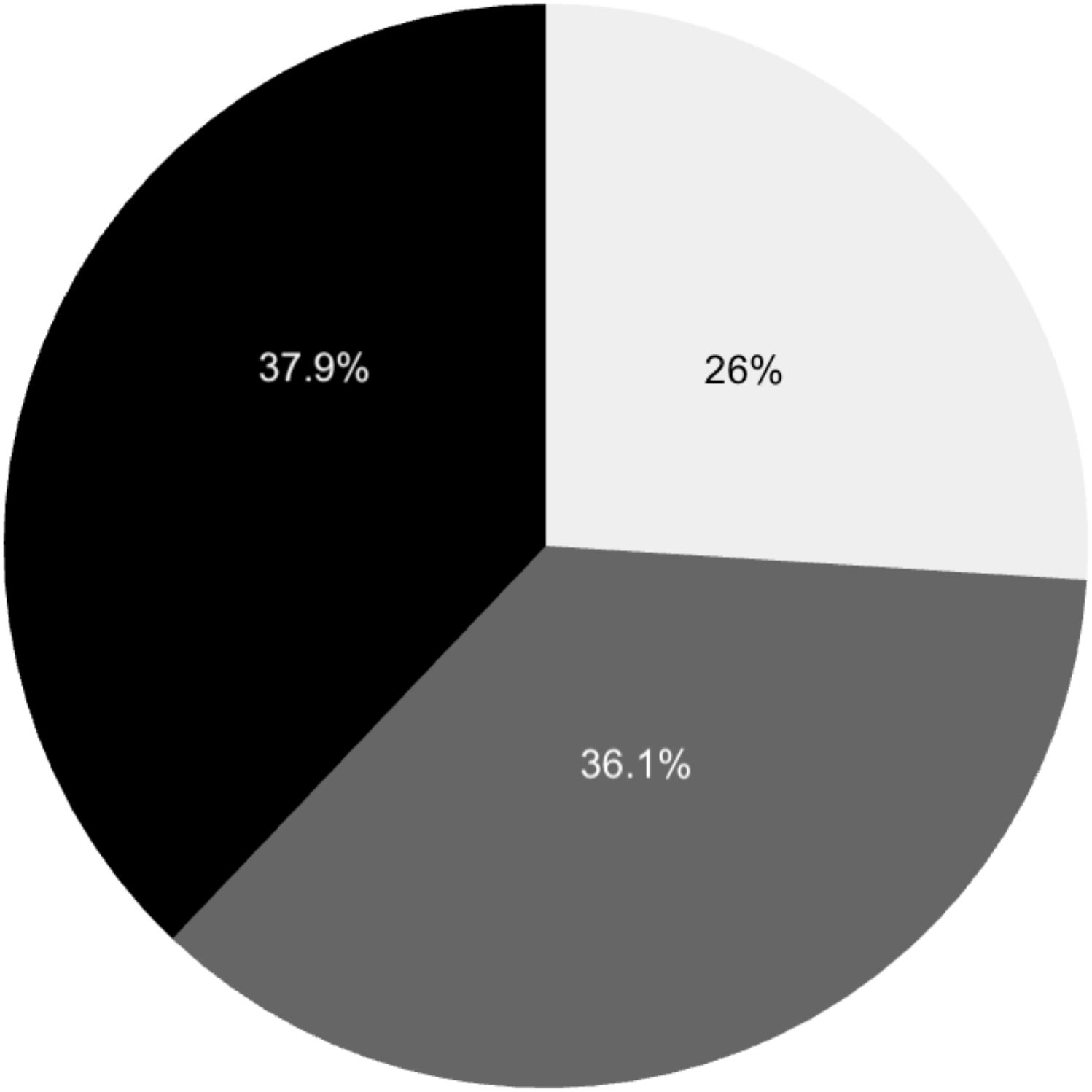
Percentage of conserved Embyrophyta genes present in *T. discolor* as identified by BUSCO analysis. Light gray - complete genes; dark gray - fragmented genes; black - missing genes.

### Candidate gene analysis

Interactions between plants and endophytic fungi are likely mediated in part by specialized metabolites (Chen and Jiang, 2026). Likewise, the production of plant specialized metabolites can be influenced by endophytic fungi (as well as other biotic and abiotic interactions). In an effort to develop a broad view of the metabolic potential of *T. discolor*, the identification of candidate genes was undertaken in the CCO collection. Candidate genes were selected to bridge carbon metabolism, sensing, and signaling with the channeling of carbon to pathways involved in signaling, symbiosis, and defense (Fig. 2).

**Figure 2.**
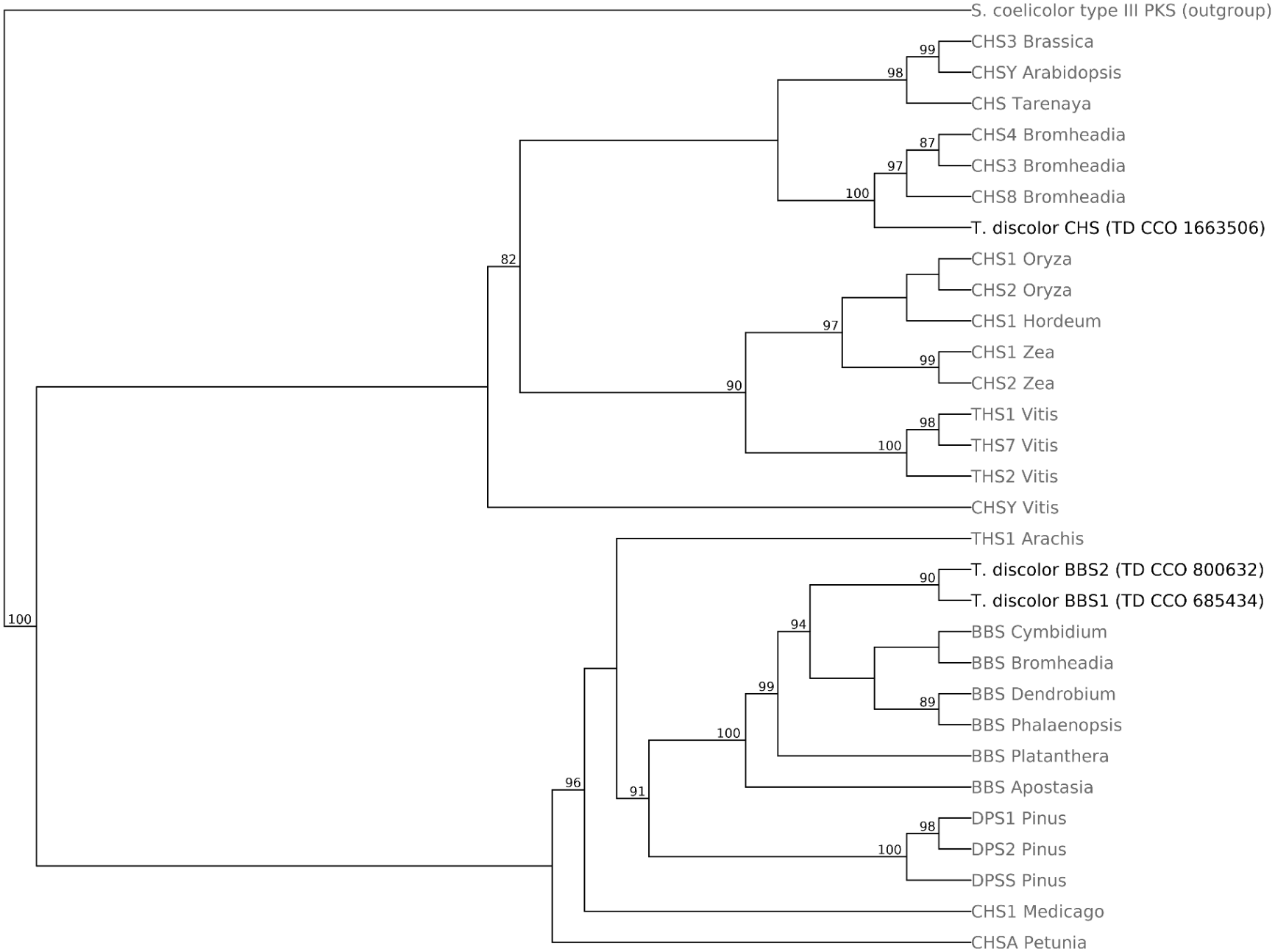
Illustrated rationale for candidate gene selection. The figure was generated in google slides and converted to PDF using Adobe Acrobat. The author’s original images were converted to cartoon using nano banana. Chemical structural line drawings were downloaded from https://www.rcsb.org/search/chemical

#### Sucrose metabolism

Candidate genes for sucrose synthase and sucrose-phosphate phosphatase were identified with strong reference-protein alignments and supporting protein-family domains (Table 5). Sucrose-phosphate synthase was identified, although domain support was not as strong. A candidate for the sucrose signaling protein SnRK1 was also identified with strong support having both correct domain architecture and conserved sequence motifs.

#### Trehalose metabolism

Candidates for both type I and type II trehalose-6-phosphate synthase (TPS) genes were identified, with strong alignments to *Arabidopsis* reference proteins and supporting protein-family domains (Table 5). A putative trehalose-6-phosphate phosphatase (TPP) candidate was also identified with high support, whereas the trehalase candidate was supported at a moderate level.

#### Phenylpropanoid metabolism

Genes associated with the mevalonate, shikimate, and flavonoid pathways were analyzed because these pathways contribute to the production of a broad array of specialized metabolites. Three enzymes of the mevalonate pathway were examined: acetyl-CoA acetyltransferase (AACT), 3-hydroxy-3-methylglutaryl-CoA synthase (HMGS), and 3-hydroxy-3-methylglutaryl-CoA reductase (HMGR). Miniprot identified partial candidate models for all three genes. Despite incomplete reference protein coverage, the identities of the HMGS and HMGR candidates were strongly supported by amino acid identity and similarity across the aligned regions and by the presence of expected Pfam domains (Table 5). The AACT candidate encoded only an N-terminal portion of the reference protein but contained a supported N-terminal thiolase domain and was therefore annotated more conservatively as an AACT-like fragment.

For the shikimate pathway, candidates for genes encoding seven enzymes were examined: 3-deoxy-D-arabino-heptulosonate 7-phosphate synthase (DAHPS), 3-dehydroquinate synthase (DHQS), 3-dehydroquinate dehydratase (DHQD), shikimate dehydrogenase (SDH), shikimate kinase (SK), 5-enolpyruvylshikimate-3-phosphate synthase (EPSPS), and chorismate mutase (CM). DAHPS and EPSPS were represented by nearly complete homologs with high identity to reference proteins and were assigned putative annotations with high confidence (Table 5). DHQS showed a strong reference-protein alignment with support for both expected N- and C-terminal domains, although the predicted model covered only 38% of the reference protein, indicating that the candidate was incomplete. A single candidate for the bifunctional DHQD/SDH protein was identified. Although reference protein coverage was incomplete, the predicted model retained both expected Pfam domains. However, the model contained three internal stop codons and was retained with only moderate confidence. Similarly, the CM candidate aligned to only a portion of the reference protein and contained an internal stop codon. SK mapped to two different CCOs, both representing fragments of full-length SK proteins, and these candidates were therefore assigned moderate confidence.

Malonyl-CoA production largely depends on conversion of acetyl-CoA by acetyl-CoA carboxylase (ACC), a key step in fatty acid biosynthesis as well as phenylpropanoid metabolism (de Souza, 2020). The initial miniprot screen identified one CCO containing a portion of an ACC gene. Targeted searches using partial ACC1 reference sequences identified three additional CCOs corresponding to different regions of the ACC1 protein (Table 5). Each candidate showed strong sequence similarity to the corresponding reference region and was supported by expected protein domains. Together, the four CCOs represent different portions of ACC1, although it cannot be determined from the fragmented assembly whether they derive from a single gene or from related members of the same gene family. Examination of sequences flanking the ACC1 candidate regions found no significant matches to known repetitive elements using RepeatMasker. However, comparison against the CCO collection identified a flanking sequence that matched 465 non-self CCOs, suggesting the presence of a highly repetitive element not represented in the repeat database.

Type III polyketide synthases catalyze the condensation of malonyl-CoA with CoA-ester starter molecules to produce a diverse array of specialized metabolites. The initial focus was on chalcone synthase (CHS), which together with chalcone isomerase (CHI) mediates entry into the flavonoid pathway. One partial CHS candidate was identified, with most of the gene represented on a single CCO but the region encoding the N-terminus absent (Table 5). A CHI candidate was also identified but was incomplete and showed lower reference-protein identity and similarity, resulting in a moderate-confidence assignment. Intriguingly, targeted searches intended to recover the missing N-terminal region of CHS identified two additional type III PKS candidates with strong similarity to orchid bibenzyl synthases (BBS). Both candidates contained the expected type III PKS domain and were represented by substantially complete protein models. Phylogenetic analysis placed the two *T. discolor* candidates together within the orchid BBS group, with 90% support for the *T. discolor* pair and 94% support for their placement with the orchid BBS sequences (Fig. 3). These candidates were therefore assigned as putative BBS1 and BBS2.

**Figure 3.**
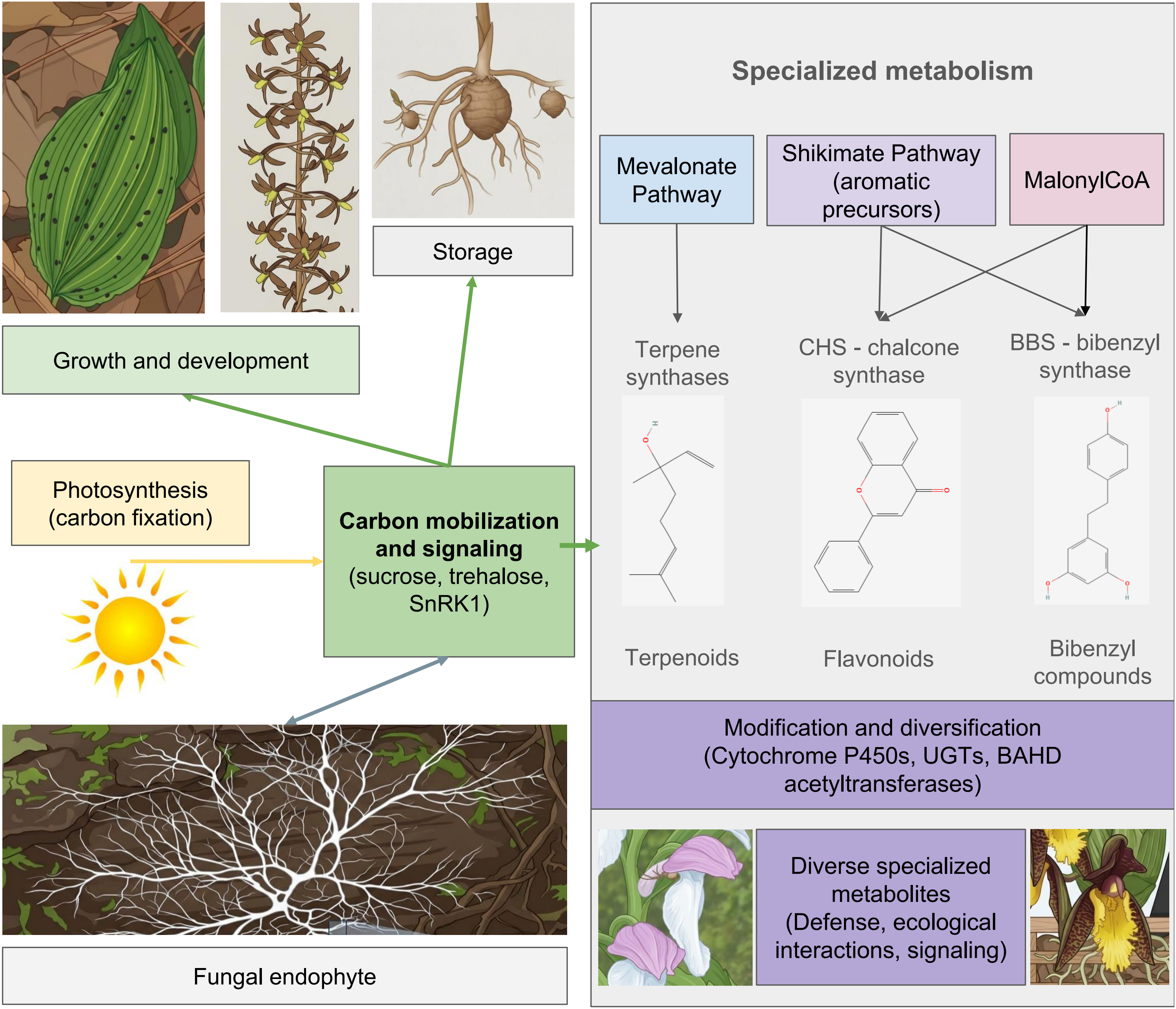
Cladogram of representative type III PKS homologs. Type III PKS candidates from *T. discolor* (black) were aligned with CHS and stilbene synthases from monocots and dicots (gray). BBS proteins from other orchids were included as were close dicot matches to BBS from *Tarenaya* and *Brassica*. All protein sequences were aligned using ClustalW and the tree generated using the fasttree option (https://www.genome.jp/tools-bin/clustalw). The Newick tree was rooted with a bacterial type III PKS from *Streptomyces coelicolor* (WP_011027653) and the subsequent tree built using ape in R. Branch lengths in the displayed cladogram do not represent evolutionary distance, and node support values ≥80% are shown. Protein genebank accession numbers are listed in Supplemental Table S5.

#### Terpene metabolism

Terpenes are found in all land plants, synthesized from two 5-carbon building blocks, isopentenyl diphosphate and dimethylallyl diphosphate, by two pathways located in distinct subcellular locations (Zhou & Pichersky, 2020) Terpene synthases (TPS) are key enzymes in both pathways, defining the backbones of the mono-, sesqui-, and diterpenes Five TPS gene candidates were identified within the CCO collection and supported by reference-protein alignments, expected Pfam domains, model completeness, and the absence of internal stop codons (Table 5). The candidates showed similarity to TPS with differing reported activities, suggesting representation of multiple TPS classes within the *T. discolor* assembly.

#### Specialized metabolite modification

Further synthesis and modification of terpenoids and other specialized metabolites can involve cytochrome P450 monooxygenases, UDP-glycosyltransferases (UGTs), and BAHD acyltransferases. Numerous initial candidates from each of these large gene families were identified within the CCO collection, and a subset with the strongest combined alignment, domain, motif, and model-completeness support was retained for further analysis (Table 5). For example, 46 initial UGT candidates were identified, of which 38 contained significant matches to the expected Pfam domain; 15 were ultimately retained based on combined reference-protein alignment, model integrity, domain, and motif evidence. Collectively, the retained P450, UGT, and BAHD candidates provide evidence for representation of diverse specialized-metabolism gene families within the *T. discolor* assembly.

Despite the fragmented nature of the assembly, candidates representing carbon metabolism, signaling, and several major specialized metabolism gene families were recovered with varying levels of support (Table 5). The strongest candidates were supported by multiple independent lines of evidence, whereas those rated with lower confidence often reflected partial or fragmented candidates.

## Discussion

*T. discolor* is a terrestrial North American orchid in the Epidendroideae subfamily that offers the potential to serve as a model for diverse aspects of orchid biology. Its position within a clade consisting largely of epiphytic members provides an opportunity to investigate mechanisms associated with terrestrial habitat utilization and adaptation. Its unique habitats relative to other orchids offers promise for expanding this knowledge and the underlying genetic components to inform conservation efforts as well as targeted crop improvement programs. Its relationship to terrestrial orchid species from Asia means that genomic analysis of *T. discolor* expands the phylogenetic and geographic range of comparative genome analyses for one of the largest and most diverse plant families. The long-term goal of this work is to establish a complete genome sequence for *T. discolor*, building on the foundational estimation of the genomic parameters determined in the current study. However, the immediate objectives were to develop a draft genomic resource with sufficient representation of gene space to identify and investigate genes relevant to orchid growth, development, and interactions with endophytic fungi.

The selection of candidate genes was directed in part by an interest in the interaction of orchids with endophytic fungi(Fig. 2). Carbohydrate flux is important to the formation and maintenance of both endo- and ectomycorrhizal symbioses and can influence the balance between symbiosis and pathogenicity (Nehls et al., 2010; Wang and Wu, 2023). Genes associated with carbon mobilization and signaling were therefore selected because they provide targets for future functional studies of *T. discolor* responses to interactions with endophytic fungi. Despite the fragmented nature of the assembly, the conservative gene-mining approach recovered candidates representing each of the targeted aspects of carbon metabolism and signaling. As expected for a photosynthetic plant, candidates associated with sucrose and trehalose metabolism as well as the carbon-signaling kinase SnRK1 were identified. Trehalose metabolism is particularly interesting in this context because trehalose and trehalose signaling have been associated with both abiotic stress and mycorrhizal interactions (Bao et al., 2024; John et al., 2017; Liu et al., 2006; Paul et al., 2008; Ponert et al., 2021). Likewise, sucrose metabolism and signaling are involved broadly in plant carbon partitioning, growth, development, and stress responses (Bagnato et al., 2023; Li et al., 2024).

Carbon mobilization is also associated with the allocation of carbon into specialized metabolic pathways involved in defense, signaling, and interactions with other organisms (Fig. 2). Candidate-gene analysis was therefore extended from sucrose and trehalose metabolism to a range of candidates associated with specialized metabolite production and modification. The recovery of said candidates demonstrates that the assembly contains gene sequences representing diverse components of specialized metabolism as has been demonstrated in the published orchid genomes(Cai et al., 2014, Zhang et al., 2016). Importantly, the genes identified here now provide targets for functional characterization and, more specifically, investigation of their role in regulating interactions with endophytic fungi.

One of the most striking outcomes of this study was the identification of two candidates encoding putative bibenzyl synthases (BBS). Bibenzyl compounds occur in several orchids used medicinally in Asia and have attracted interest because of their reported pharmaceutical activities (Boddington et al., 2022; Li et al., 2025; Liu et al., 2024; Patel et al., 2024). The two candidates were initially recovered during targeted searches for the missing N-terminal region of the partial *T. discolor* CHS candidate. Although both encoded CHS-like type III PKSs, sequence comparisons showed stronger similarity to orchid BBS proteins. Phylogenetic analysis further supported this assignment: the two *T. discolor* proteins clustered together with 90% support and joined the orchid BBS group with 94% support (Fig. 3). In contrast, the partial *T. discolor* CHS candidate clustered with reference CHS proteins, supporting the distinction between the CHS and BBS candidates. The presence of these putative BBS homologs raises the possibility that bibenzyl or bibenzyl-derived metabolites occur in *T. discolor*, although enzymatic activity and the presence of corresponding metabolites remain to be demonstrated.

In summary, analysis of the *T. discolor* genome using Illumina short-read sequencing indicated a relatively large genome of approximately 1.7 Gb with a high proportion of repetitive sequence. Both ABySS and MaSuRCA generated highly fragmented assemblies, likely reflecting the difficulty of assembling repetitive regions from short reads. However, the MaSuRCA CCO collection contained 62.1% of conserved Embryophyta BUSCOs as either complete or fragmented sequences, indicating substantial representation of conserved gene space. Targeted mining of this resource identified 68 candidate genes associated with carbon metabolism and signaling as well as various aspects of specialized metabolism. These candidates are now targets for functional analysis and demonstrate that, despite its fragmentation, the current assembly can support targeted discovery of genes associated with specific biological questions. While the current assembly cannot reliably resolve gene copy number, genomic organization, promoter structure, gene clusters, or chromosome-level synteny. However, it provides a genomic resource that can be further mined for additional genes of interest. Furthermore, it can support the development of future expression studies, including, for example, RNA-seq analyses of *T. discolor* responses to fungal and other environmental interactions. Future long-read sequencing will be required to generate a more complete genome assembly and provide the genomic context unavailable from the present short-read resource.

Taken together, these continuing efforts will provide a foundation for investigating the molecular mechanisms underlying the growth, development, specialized metabolism, and ecological interactions of *T. discolor*.

## Acknowledgements

The author acknowledges the support of Brena Winkel for reading and critical analysis of the manuscript. This work was supported through annual internal teaching funds.

## Author contributions

Jonathan Watkinson performed all analyses and wrote the manuscript.

